# Effects of substrate stiffness and actin velocity on *in silico* fibronectin fibril morphometry and mechanics

**DOI:** 10.1101/2021.02.24.432650

**Authors:** Seth H. Weinberg, Navpreet Saini, Christopher A. Lemmon

## Abstract

Assembly of the extracellular matrix protein fibronectin (FN) into insoluble, viscoelastic fibrils is a critical step during embryonic development and wound healing; misregulation of FN fibril assembly has been implicated in many diseases, including fibrotic diseases and cancer. We have previously developed a computational model of FN fibril assembly that recapitulates the morphometry and mechanics of cell-derived FN fibrils. Here we use this model to probe two important questions: how is FN fibril formation affected by the contractile phenotype of the cell, and how is FN fibril formation affected by the stiffness of the surrounding tissue? We show that FN fibril formation depends strongly on the contractile phenotype of the cell, but only weakly on tissue stiffness. These results are consistent with previous experimental data and provide a better insight into conditions that promote FN fibril assembly. We have also used the model to look at two distinct phenotypes of FN fibrils that we have previously identified; we show that the ratio of the two phenotypes depends on both tissue stiffness and contractile phenotype, with intermediate contractility and high tissue stiffness creating an optimal condition for stably stretched fibrils. Finally, we have investigated how re-stretch of a fibril affects cellular response. We probed how the contractile phenotype of the re-stretching cell affects the mechanics of the fibril; results indicate that the number of myosin motors only weakly affects the cellular response, but increasing actin velocity results in a decrease in the apparent stiffness of the fibril and a decrease in the stably-applied force to the fibril. Taken together, these results give novel insights into the combinatorial effects of tissue stiffness and cell contractility on FN fibril assembly.

## Introduction

### Fibronectin plays a prominent role in embryonic development, wound healing, and disease

Fibronectin (FN) fibrils are the primordial extracellular matrix assembled by fibroblasts during wound healing and embryogenesis. In embryogenesis, FN fibrils are essential for some of the earliest developmental steps: in Xenopus embryos, gastrulation fails in the absence of FN, and the embryos have significant cardiovascular defects [1]. In wound healing, the initial fibrin clot binds to factor XIII which in turn facilitates binding with FN [2]. FN fibrils serve both as a means of structurally stabilizing the clot [3] and as a network that facilitates cell migration into the wound to direct immune response and tissue building [4].

### Cells stretch fibronectin to drive assembly into insoluble, viscoelastic fibrils

Fibronectin is a 250-kDa glycoprotein that is present in a soluble form at high concentration in the blood plasma [5]. Cells bind to soluble plasma fibronectin via transmembrane integrins. Integrins are stretched via contraction of the actomyosin cytoskeleton, which transmits force via focal adhesions to integrins and then to FN [6]. Stretching FN exposes buried cryptic FN-FN binding sites, which leads to the incorporation of another soluble plasma FN [7]. This in turn leads to assembly of networks of viscoelastic, insoluble fibrils.

### Assembly of fibronectin fibrils is dependent on cell phenotype

Since FN fibrils require cell-derived traction forces to expose cryptic binding sites [8], and since different cell types generate differing magnitudes of contractile forces [9], it thus follows logically that the assembly of FN fibrils depends on the cell type. Previous studies have demonstrated that fibroblast and mesenchymal cells robustly assemble fibrils in vitro, while epithelial and endothelial cells exhibit negligible fibrillogenesis [9]. There is a bi-phasic response however; cells that generate the largest forces, such as smooth muscle cells and cardiomyocytes, also generate only minimal numbers of fibrils [9]. One way in which we can represent these cellular differences in contractile state is by varying actin velocity, specifically the unloaded velocity of an actin filament not tethered to a focal adhesion. Prior studies have demonstrated that this unloaded actin velocity varies over several orders of magnitude between differing cell types [10]. These studies show that cells with higher unloaded actin velocities correspond with cells that have been shown to exhibit higher contractility.

### Assembled fibronectin fibrils alter the mechanoresponse of attached cells

Over the past two decades, there has been an increased understanding of the role that tissue stiffness plays in driving cellular mechanoresponses [11–13]. It is now well appreciated that cells on increasingly stiffer surfaces generate larger traction forces. FN fibrils play a key role in the sensing of substrate stiffness; when FN fibril assembly is inhibited, fibroblasts do not generate larger forces in response to increasing tissue stiffness [14]. Despite this key role in mechanosensing, there does not seem to be a correlation between substrate stiffness and the degree of FN assembly: previous studies from our group have demonstrated that cells will assemble FN fibrils across a range of substrate stiffness values, with only a slight dependence on substrate stiffness [14].

### A computational model of fibronectin fibril formation captures in vitro fibril morphometry and mechanics

The mechanism of FN fibril assembly is still poorly understood. This is partially due to the fact that, as opposed to actin or tubulin which spontaneously assemble in cell-free environments, FN fibril assembly requires cell-generated forces, and is thus difficult to replicate in cell-free environments. While several groups have attempted to generate FN fibrils in cell-free environments by shearing soluble FN [15–17], these fibrils often exhibit different morphologies and mechanics from cell-derived FN fibrils. To probe the potential mechanism of FN fibril mechanics, we previously generated a computational model that constructs *in silico* FN fibrils from first principles [18]. This model was based on a previously developed motor-clutch model that simulated the interaction between the actomyosin contractile unit of the cell and the underlying substrate [19] and predicted cellular mechanical responses to various substrate stiffnesses. In our previous work, we integrated a fibronectin molecule between the cell and substrate, and then incorporated the mechanism of FN incorporation and fibril growth (shown in Fig. 1, described further in Materials and Methods, and described fully in [18]). We then compared the morphology and mechanics of these simulated fibrils to cell-derived fibrils, and have shown that the simulated fibrils recapitulate many of the fibril properties, including fibril length, length-to-thickness ratio, and stretched-to-relaxed length ratio.

**Fig 1.**
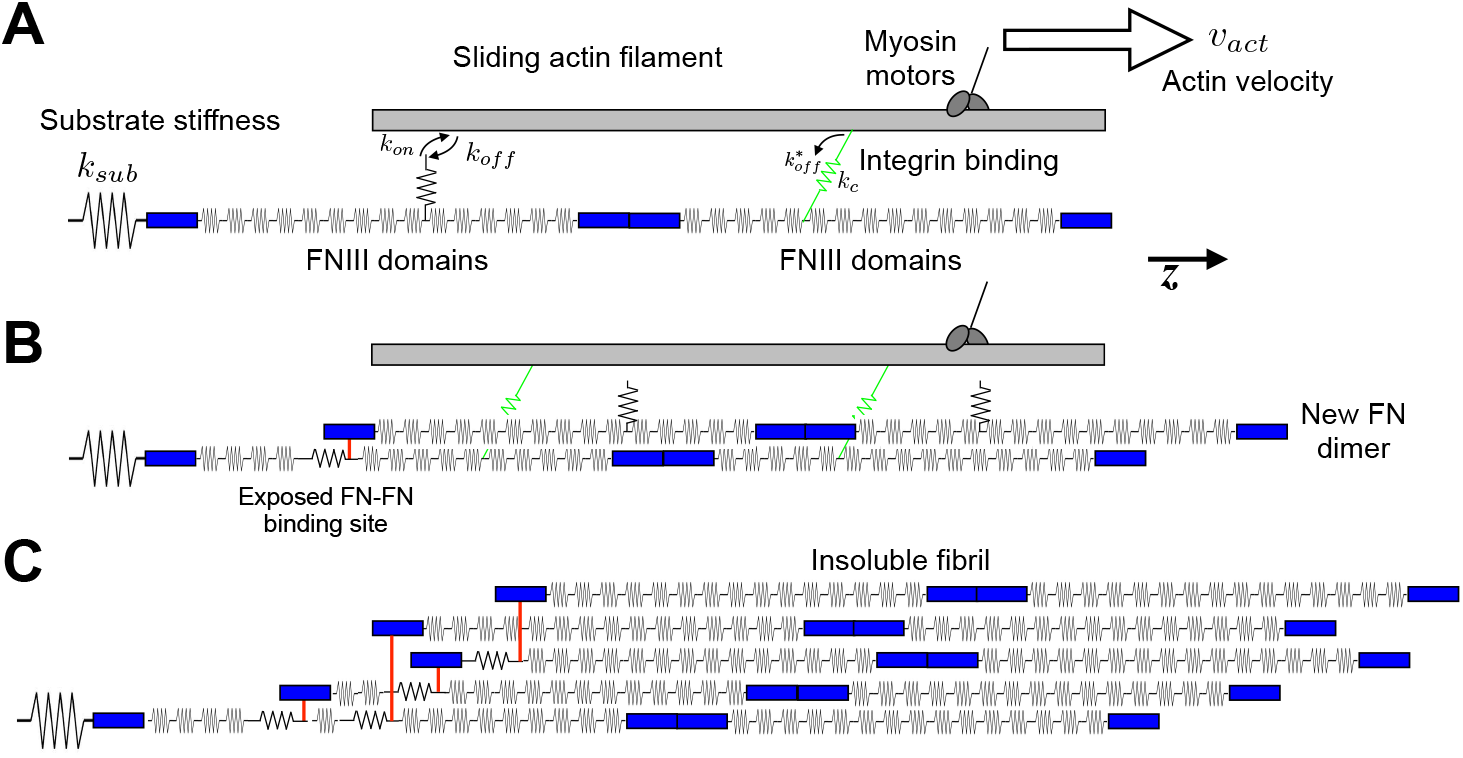
Diagram of computational model of FN fibril assembly. A schematic of the FN fibril assembly model used in the current study. A) Actomyosin machinery stochastically binds to FN at the III-10 domain, with a force-dependent *k*_*off*_. B) This binding and pulling unfolds FN Type III domains, exposing cryptic FN-FN binding sites. C) This process continues until the system reaches equilibrium, such that the fibril, clutch and mysoin pulling force are balanced, and no new FN molecules are added.

Previous studies suggest a potentially complex relationship between cell phenotype, tissue stiffness, and FN fibril formation. In the current work, we investigate these relationships by using our existing FN fibril assembly model to investigate the following questions: first, how does substrate stiffness affect FN fibril formation in our model? Our prior experimental work would suggest that the model should predict FN fibril assembly across a range of stiffness values. Second, how does cellular contractile phenotype, as characterized by the unloaded actin velocity, affect FN fibril assembly? Prior studies from our group and others have suggested that the contractile phenotype of the cell strongly affects FN fibril assembly, regardless of substrate stiffness. Finally, how does the contractile phenotype of a cell that is re-stretching a previously assembled fibril affect its mechanosensing response? In other words, if two cells of differing contractile phenotype migrate over an existing FN fibril and re-stretch it, do the cells “feel” a different fibril? By answering these questions, we can gather significant insight into the mechanism of FN fibril assembly in response to a range of mechanical environments and cellular phenotypes.

## Materials and methods

### Model Formation

To simulate fibronectin fibril growth, we used a hybrid stochastic-deterministic model that we have previously developed [18]. A schematic of the model is shown in Fig 1. Briefly, we model an actin filament, pulled on by a myosin motor and coupled to a “clutch”, which represents the collective focal adhesion/integrin complex. This clutch is attached to a single FN molecule, which in turn is coupled to the underlying tissue. The myosin motor pulling on the actin filament is modeled using the Hill force-velocity relationship, in which the resistive force is inversely proportional to the actin velocity:

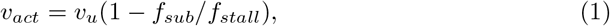

where *f*_*sub*_ is the force acting on the substrate, *f*_*stall*_ = *N*_*myo*_ *f*_*myo*_ is a myosin stall force for which *v*_*act*_ approaches 0 and that is proportional to the number of myosin motors *N*_*myo*_ and individual myosin stall force *f*_*myo*_, and *v*_*u*_ is the unloaded actin velocity.

The focal adhesion “clutch” is represented by Hookean springs in parallel, each of which has a force-dependent unbinding rate, which drives integrin/FN rupture proportional to force generation. The FN molecule is represented by 30 springs in series, which represent the elastic Type III domains in FN, which have previously been shown to unfold in response to force [20]. Finally, the tissue/substrate is modeled as a single Hookean spring, with substrate stiffness *k*_*sub*_.

Simulations begin initially with a single (unbound) FN molecule, and the actin filament moving at its unloaded actin velocity *v*_*u*_, a parameter of the simulation. The springs that comprise the clutch stochastically bind and unbind, attaching to the FN molecule, which in turn transmits force to the spring representing the substrate. The resultant motion generates a reaction force. This reaction force alters both the speed of the actin filament and the binding/unbinding kinetics of the focal adhesion clutch. The force also unfolds Type III domains in FN; once a domain has exceeded a length threshold (determined based on published steered molecular dynamics simulations of Type III domains [21]), it is deemed “open” and can bind a second FN molecule. This new molecule creates new binding sites for the focal adhesion/integrin clutch springs. The simulation proceeds until assembly is completed: that is, the fibril, clutch, and myosin pulling force are balanced such that no new addition of FN molecules occurs. In order to model the 3-dimensional aspect of an FN fibril, new FN molecules are added to the existing fibril using a hexagonal geometry. New FN molecules stochastically fill an empty site in the hexagonal pattern, and only FN molecules on the “outside” of the fibril are able to bind new FN molecules or integrin clutches. An in-depth discussion of the computational algorithm, including the handling of deterministic-stochastic coupling, can be found in the original publication of the model [18].

In this study, we have conducted new “re-stretching” simulations of *in silico* FN fibrils as follows: the result of the simulations above is a stretched FN fibril, with many Type III domains stretched open and binding to other FN molecules. To probe the mechanics of the FN fibrils, we “relax” these fibrils by resetting all FN Type III springs and the substrate spring to their resting lengths. This fibril is “re-stretched” by simulating the same myosin force, actin filament speed, and focal adhesion clutch binding as in the previous assembly simulations. Importantly, these parameters can differ from those used in the original simulation, allowing us to simulate a fibril assembled with one set of actomyosin parameters, but re-stretched with a different set of parameters. Note that in these re-stretching experiments, the addition of new FN molecules is inhibited.

Our prior studies have demonstrated two distinct FN fibril “phenotypes”: one in which the simulation stabilizes at a constant FN fibril length with a constant applied force, which we have termed “Stably Stretched Fibrils (SSFs)”; and one in which the simulation never stabilizes, but stochastically fluctuates around a fibril length and applied force, which we have termed “Fluctuating Stretched Fibrils (FSFs)”. Fig. 2A shows a representative displacement vs. time plot for a fibril of each phenotype, Fig. 2B shows a representative force vs. time plot for a representative fibril from each phenotype, and Fig. 2C shows a representative force versus stretch plot for a representative fibril from each phenotype. An in-depth discussion of these two FN fibril phenotypes can be found in our prior work [22].

**Fig 2.**
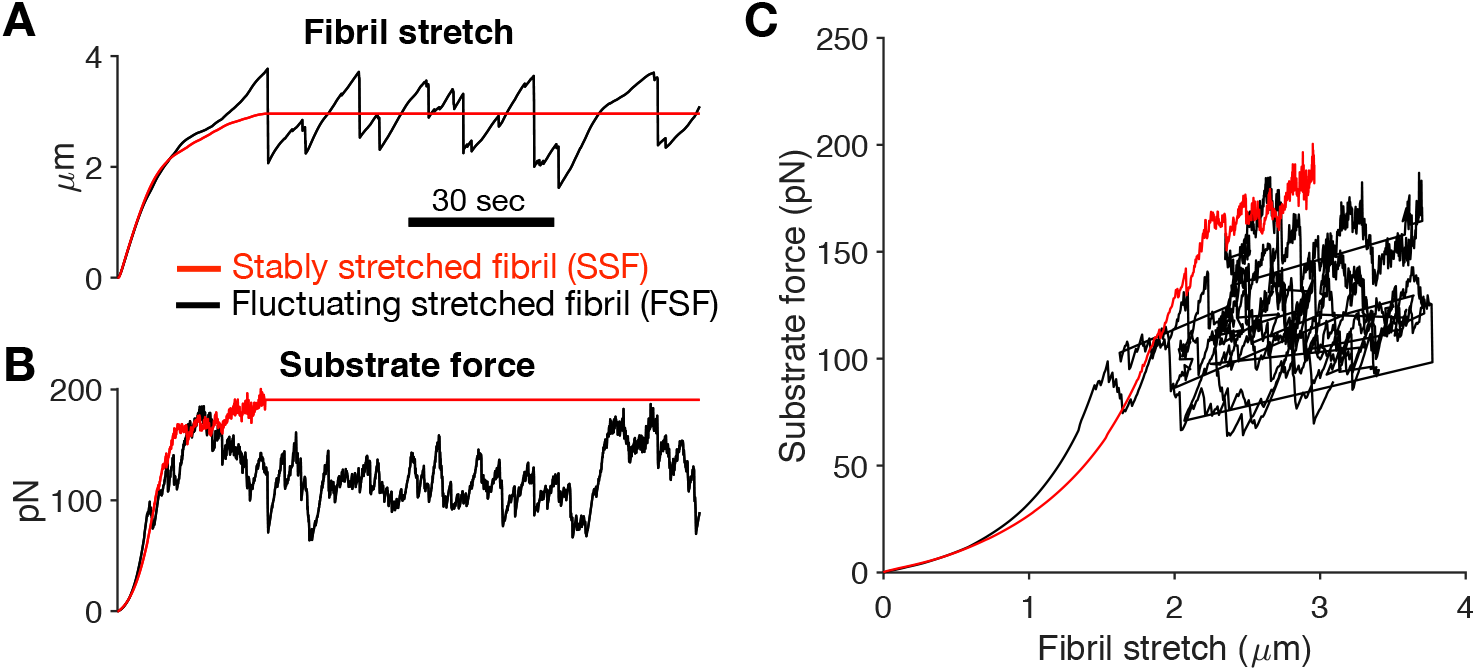
Stably stretched FN fibrils versus fluctuating stretched FN fibrils. *In silico* FN fibrils exhibit two distinct phenotypes. A) SSFs reach a stable stretch length, while FSF fluctuate around a central value; B) SSFs exhibit constant force transmission to the underlying substrate, while FSFs fluctuate; C) Force/dsiplacement curves for a representative SSF (red) and FSF (black).

## Model Simulations

In the current work, we investigate two parameters: 1) the stiffness of the Hookean spring is changed to represent tissues of varying stiffness *k*_*sub*_; 2) the unloaded actin velocity v_*u*_, which is the speed at which myosin pulls the actin filament in the absence of any resistive force, is changed to represent different contractile phenotypes. A higher unloaded actin velocity represents a larger pulling force by mysoin; previous experimental studies have demonstrated that unloaded actin velocity varies by several orders of magnitude from approximately 7 to 5000 nm/s, depending on cell type [10]. In a subset of experiments, the number of myosin motors *N*_*myo*_ was varied. For each combination of parameters, 100 simulations were run.

## Results

### Effects of tissue stiffness and actin velocity on FN fibril morphometry

We first investigated the effects of tissue stiffness and cellular phenotype by varying the stiffness of the spring that represents the underlying substrate (*k*_*sub*_) and the unloaded actin velocity of the pulling myosin motor (*v*_*u*_), where a higher value represents a cell with a stronger contractile phenotype. Results from these simulations are shown in Fig. Mean values at the conclusion of the simulation are shown for each of the 100 simulations per condition for the following metrics: the total number of FN molecules that have been incorporated into the fibril (Fig. 3A); the total force transmitted to the underlying substrate (Fig. 3B); the length-to-thickness ratio of the fibril, which quantifies the geometry of the fibril (Fig. 3C); the stretched length of the fibril, which represents the length while the fibril is still under cell tension (Fig. 3D); the relaxed length, which represents the fibril length after release from cell tension (Fig. 3E); the ratio of these two lengths, which indicates the extensibility of the fibril (Fig. 3F); the fraction of clutches bound, which can be thought of as a corollary to the degree of integrin attachment to the fibril (Fig. 3G); the length of the region from first attachment to last attachment to the fibril, which can be thought of as a corollary to focal adhesion length (Fig. 3H); and finally, the time in hours until the simulated fibril reaches a steady state length (Fig. 3I). All reported values show a strong correlation with the unloaded actin velocity v_*u*_, but little to no dependence on tissue stiffness. The one exception is the FN fibril assembly time, which shows negligible dependence on either substrate stiffness or actin velocity.

**Fig 3.**
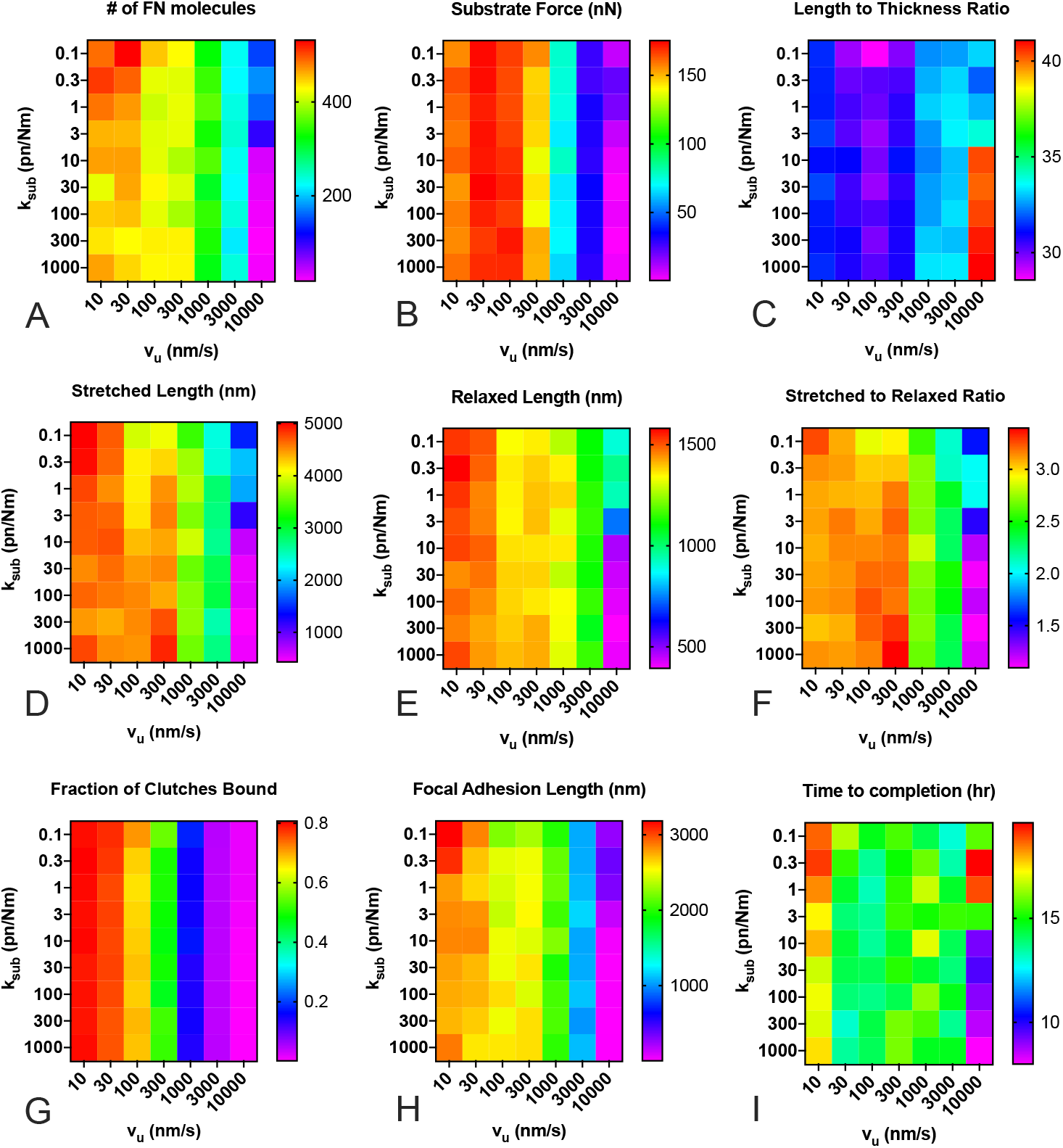
FN fibril morphometry as a function of substrate stiffness and actin velocity. Actin velocity has a pronounced effect, while substrate stiffness only shows a significant effect at extremely high actin velocities for (A) number of FN molecules in the FN fibril; (B) force transmitted via fibril to the substrate; (C) length to thickness ratio of the stretched fibril; (D) stretched length of the fibril; (E) relaxed length of the fibril; (F) stretched to relaxed ratio; (G) fraction of transmembrane clutches bound to the fibril; (H) length of focal adhesion attached to the fibril; and (I) time for FN fibril to reach steady-state. Each condition is the mean of 100 simulations.

To further analyze this response, we averaged each output in Fig. 3 for a given actin velocity, regardless of tissue stiffness (Fig. 4). Data indicate small variations at all actin velocities except for v_*u*_ = 10,000 nm/s, demonstrating that tissue stiffness does not dramatically affect FN mophometry, except at extremely high actin velocities. The data also reveal several interesting trends: as actin velocity increases, FN fibrils generated are smaller, as indicated by both the total number of FN molecules (Fig. 4A) and the total stretched and relaxed fibril length (Fig. 4D and E). As actin velocity increases, the forces transmitted through the fibril decreases (Fig. 4B), the percentage of bound integrin clutches decreases (Fig. 4G), and the average focal adhesion length decreases (Fig. 4H). These data indicate that our model is able to capture a counter-intuitive experimental finding that cells with an extremely pronounced contractile phenotype assemble shorter, thicker FN fibrils [23]. This potentially explains why cells that generate large contractile forces, such as smooth muscle cells and cardiomyocytes, exhibit negligible FN fibrilogenesis.

**Fig 4.**
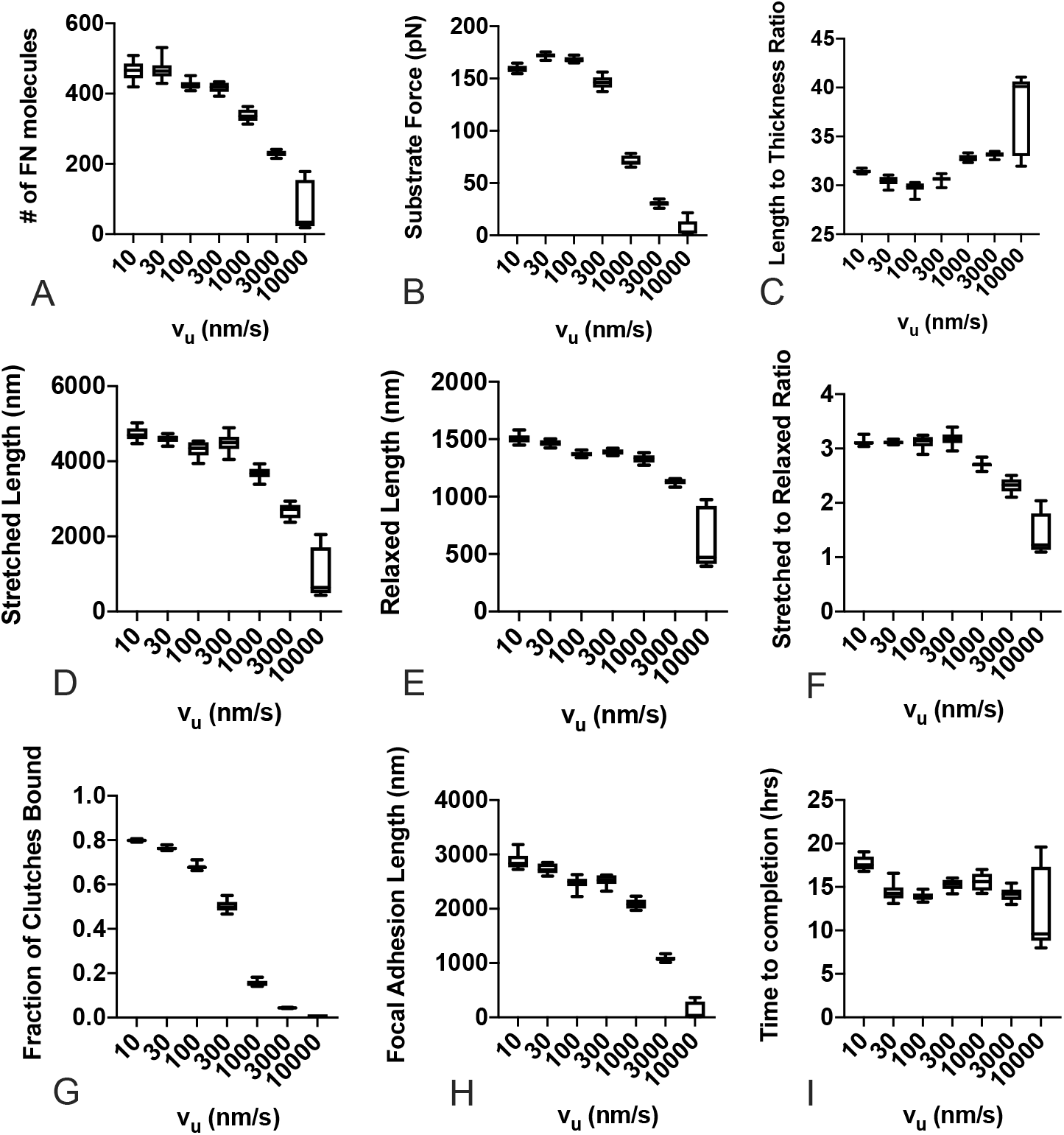
Effects of actin velocity on FN fibril morphometry and mechanics across all substrate stiffness values. Effects of unloaded actin velocity, regardless of substrate stiffness on (A) number of FN molecules in the FN fibril; (B) force transmitted via fibril to the substrate; (C) length to thickness ratio of the stretched fibril; (D) stretched length of the fibril; (E) relaxed length of the fibril; (F) stretched to relaxed ratio; (G) fraction of transmembrane clutches bound to the fibril; (H) length of focal adhesion attached to the fibril; and (I) time for FN fibril to reach steady-state. Each box and whisker point shows the mean (line), the 25th - 75th percentile range (box), and the min and max (whiskers) for 900 simulations (100 simulations for each of 9 stiffness values).

### Effects of substrate stiffness and actin velocity on FN fibril restretching mechanics

We next examined whether changing substrate stiffness or actin velocity had effects on the mechanics of FN fibrils. To investigate this, we again simulated assembly of 100 fibrils per condition as above. We then “relaxed” the fibrils by releasing tension on the FN fibril springs, resetting each spring to zero displacement. These fibrils were re-stretched at the same actin velocity that was used to assemble the fibril. Re-stretching simulations were run for up to 5 minutes, or until the fibril “stalled” at a specific stretch length and force (i.e., the actin velocity falling less than 1% of the unloaded velocity). The resulting displacement and force generation were calculated to generate a force-displacement curve for each fibril. Data from a representative re-stretch simulation is shown in Fig. 5.

**Fig 5.**
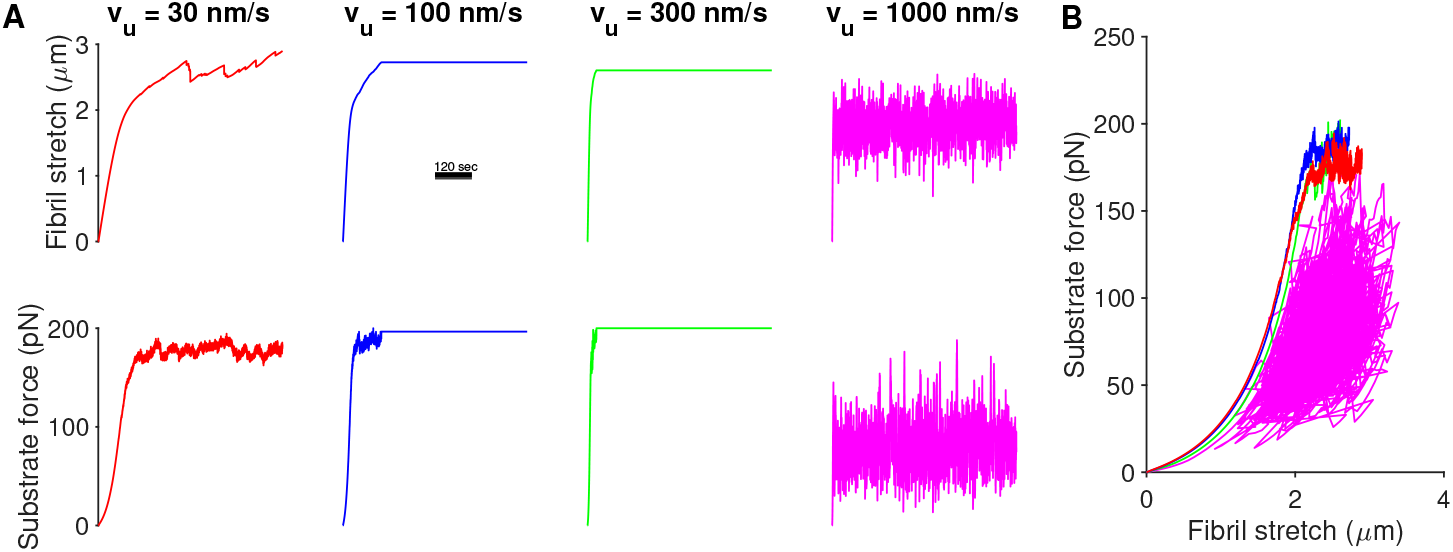
Determining the mechanical properties of FN fibrils by re-stretch simulation. *In silico* fibrils were generated as described above. After FN fibril assembly termination, fibrils were reset to their relaxed length and restretched using the same unloaded actin velocity value that was used during assembly. (A) representative stretch and force plots for 4 different unloaded actin velocities. (B) Force-displacement curves for the 4 fibrils shown in (A).

Stress in the fibril was calculated by dividing force by the maximum fibril cross sectional area, and strain was calculated as fibril stretch divided by the unstretched fibril length. The slope of the stress-strain curve gives a measurement of the fibril elastic modulus or “stiffness”. For analysis, we compared the stress-strain curve slope at the initiation of stretch. Fig. 6 shows the mean value of A) the initial slope of the force-displacement curve, B) the mean stress at the end of re-stretching, C) the mean strain at the end of re-stretch, and D) the mean fibril displacement at the end of stretching. Results indicate that substrate stiffness again has little effect, while actin velocity has a pronounced effect on FN fibril mechanics.

**Fig 6.**
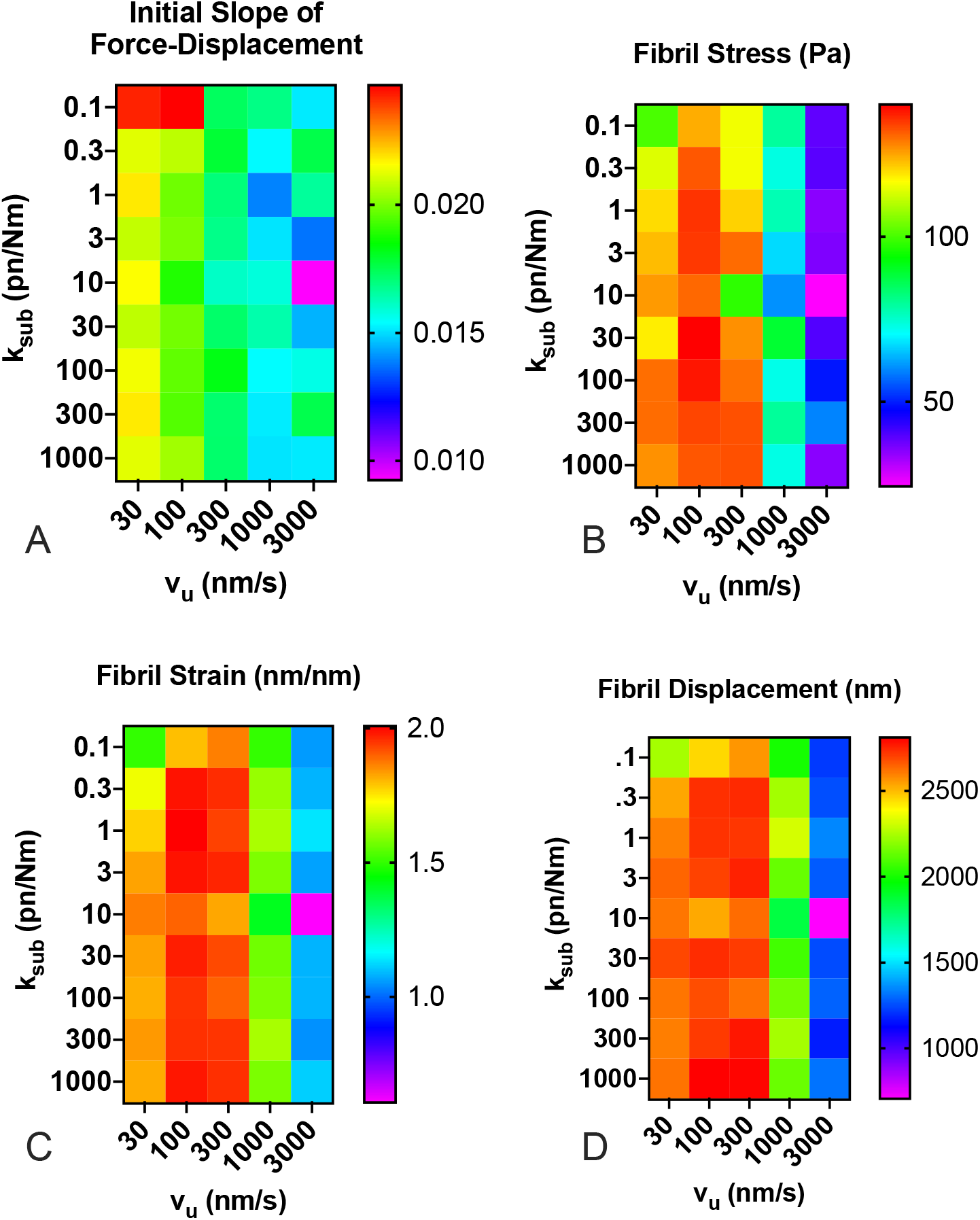
FN fibril mechanics as a function of substrate stiffness and actin velocity. Fibrils were computationally “relaxed”, and then restretched at the same actin velocity used to assemble the fibril to quantify fibril mechanics. Actin velocity had a pronounced effect, while substrate stiffness had only a moderate effect on (A) the initial slope of the force-displacement curve, (B) the fibril stress at full stretch, (C) fibril strain at full stretch, and (D) fibril displacement. Each condition is the mean of 100 simulations.

Given this predominance of actin velocity on FN fibril mechanics, we again analyzed the effects on FN fibril mechanics as solely a function of actin velocity, regardless of tissue stiffness (Fig. 7). Results indicate that the initial slope of the force-displacement curve, which indicates the initial apparent stiffness as the fibril is first stretched, decreases as actin velocity increases. This suggests that cells with a more contractile phenotype assemble less rigid fibrils, although this effect is small. Fibril stress and strain both show a maximum at intermediate values of actin velocity, suggesting that there is an unloaded actin velocity value in the 100-300 nm/s range in which fibrils exhibit a maximum stress and strain. Fibril displacement at the end of stretch only weakly depends on actin velocity, but decreases at large actin velocities.

**Fig 7.**
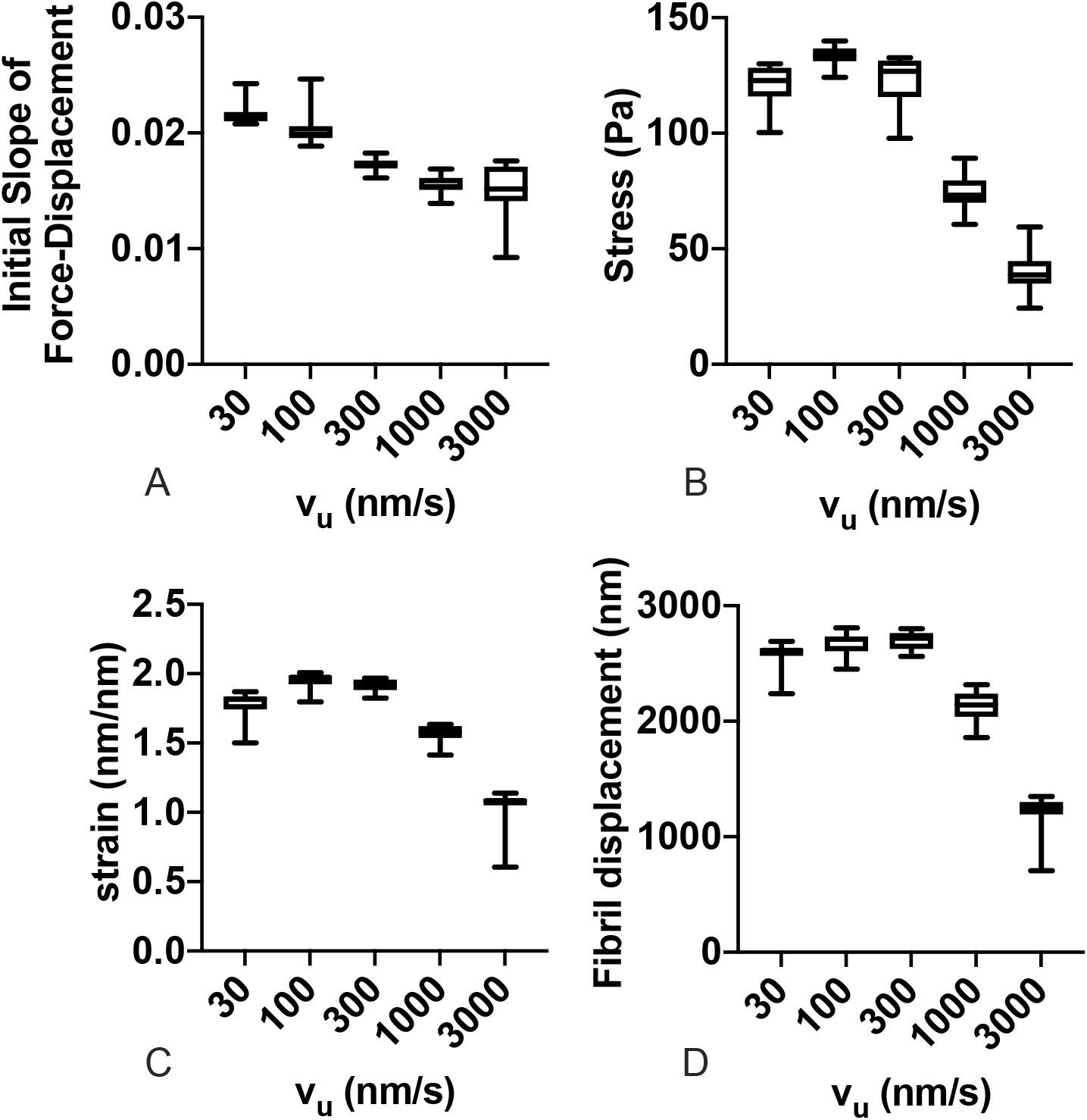
FN fibril mechanics as a function of actin velocity. Effects of unloaded actin velocity, regardless of substrate stiffness for (A) the initial slope of the force-displacement curve, (B) the fibril stress at full stretch, (C) fibril strain at full stretch, and (D) fibril displacement. Each box and whisker point shows the mean (line), the 25th - 75th percentile range (box), and the min and max (whiskers) for 900 simulations (100 simulations for each of 9 stiffness values).

### Effects of substrate stiffness and actin velocity on FN fibril phenotype

Taken together, there are three conclusions that can be drawn from the results presented so far: first, tissue stiffness has minimal effects on FN fibril morphometry and FN fibril mechanical properties; second, increasing actin velocity yields FN fibrils that are smaller and softer; and third, an intermediate unloaded actin velocity exists at which stress and strain in the fibril are maximal. We next sought to determine the effects of tissue stiffness and actin velocity on FN fibril phenotype. As discussed above, we have previously identified two distinct FN fibril phenotypes: SSFs, which remain statically stretched, and FSFs, which fluctuate around a value of force and stretch length. Fig. 8A shows the fraction of FN fibrils that exhibit the SSF phenotype. Results show a dependence on both tissue stiffness and actin velocity: at either low or high actin velocities, the phenotype is independent of tissue stiffness and is predominantly FSFs, but at intermediate actin velocities, the phenotype strongly depends on tissue stiffness: the percentage of SSFs increases with increasing tissue stiffness.

**Fig 8.**
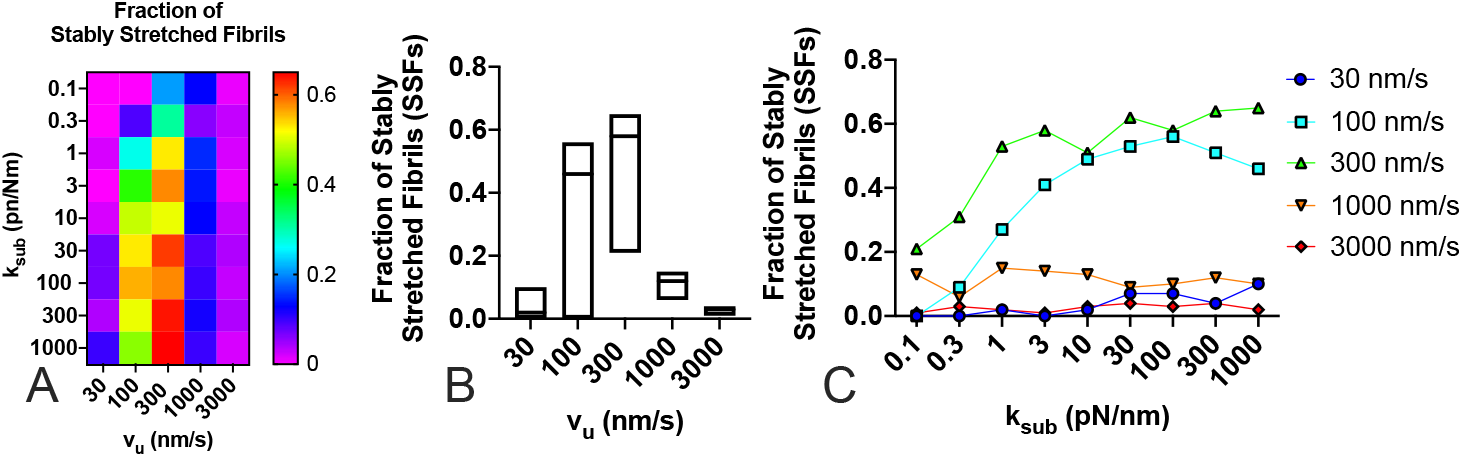
Fraction of stably stretched FN fibrils as a function of substrate stiffness and actin velocity. The fraction of fibrils that exhibited the SSF phenotype was determined for (A) both substrate stiffness and actin velocity; (B) actin velocity alone), and (C) as a function of substrate stiffness for each unloaded actin velocity.

### Effects of cell restretching properties on fibril mechanics

Cells can assemble FN fibrils, and then migrate away from them, leaving them to be re-stretched by other migrating cells. We next investigated whether the apparent properties of an assembled fibril changed depending on the contractile state of the cell. For these experiments, we used a population of 100 fibrils that were generated under the same conditions: an unloaded actin velocity of 300 nm/s, a substrate stiffness of 1000 pN/nm, and 100 myosin motors. These fibrils were released of any tension, and then res-stretched as explained above. The re-stretching of fibrils was performed over a range of values for the number of myosin motors and the unloaded actin velocity. Results, shown in Fig. 9, indicate that the apparent mechanics of the re-stretched fibril is only minimally affected by the number of myosin motors. However, the actin velocity has pronounced effects. To probe the effects of actin velocity, data was re-analyzed as a function of actin velocity only, regardless of the number of myosin motors (Fig. 10). Several interesting trends can be observed. First, there is an optimal actin velocity at which the percentage of stably stretched fibrils is maximal (Fig. 10A). This suggests that the fibril phenotype depends on both the fibril itself as well as the phenotype of the cell that is stretching it. Second, increasing actin velocity, which corresponds with a cell with an increased contractile phenotype, results in fibrils that have reduced steady state force (Fig. 10B), reduced apparent stiffness (Fig. 10C), reduced stress (Fig. 10D), reduced strain (Fig. 10E), and reduced displacement (Fig. 10F). This somewhat counter-intuitive result suggests that a cell with a stronger contractile phenotype will stretch a fibril less and exert less steady state force on the fibril compared to a cell with a lower contractile phenotype.

**Fig 9.**
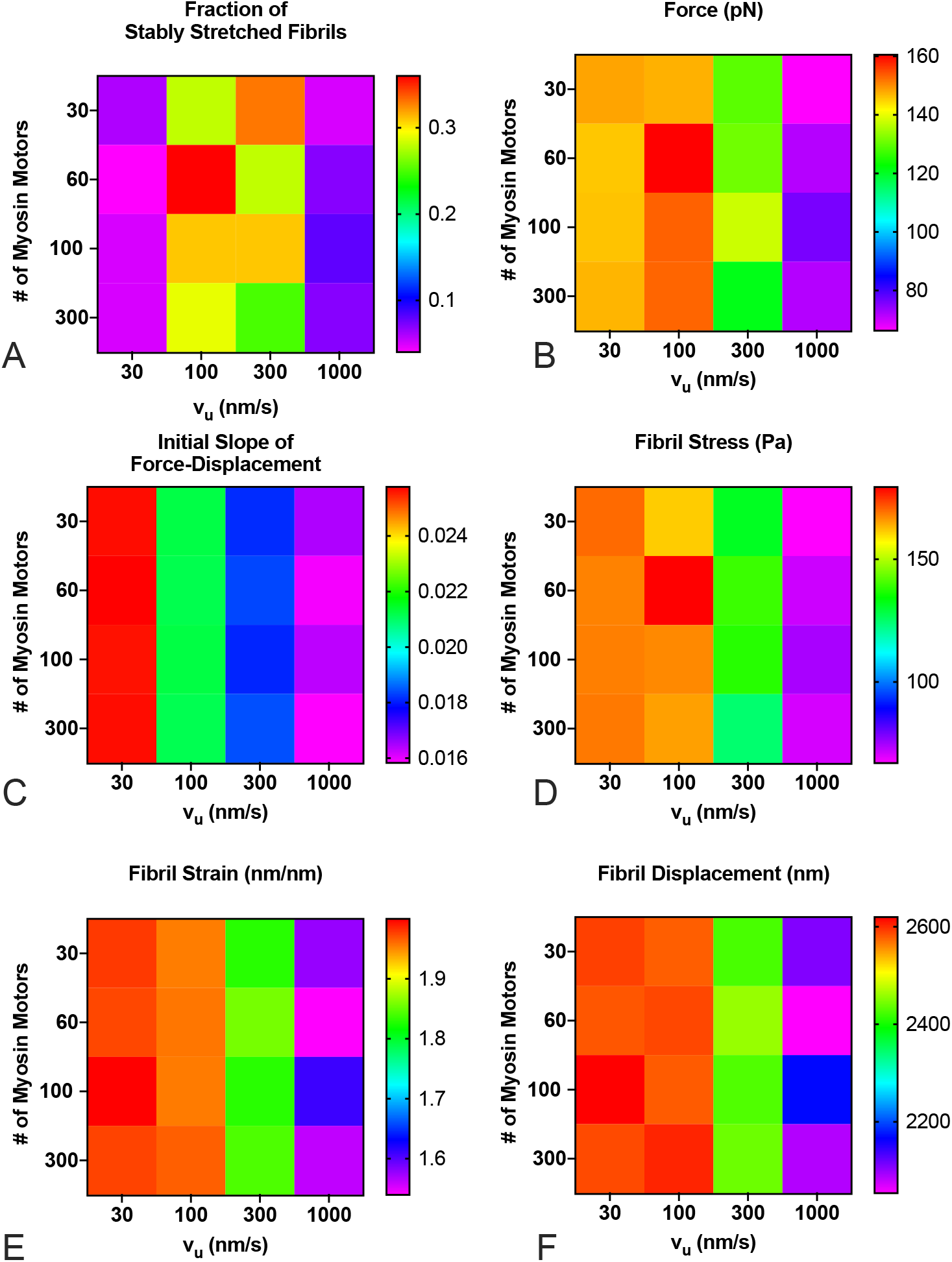
FN fibril morphometry and mechanics as a function of actin velocity and number of myosin motors. The number of myosin motors has minimal effects, while the unloaded actin velocity has significant effects on (A) the fraction of SSFs; (B) force transmitted via fibril to the substrate; (C) the initial slope of the force-displacement curve; (D) fibril stress; (E) fibril strain; and (F) fibril displacement. Each condition is the mean of 100 simulations.

**Fig 10.**
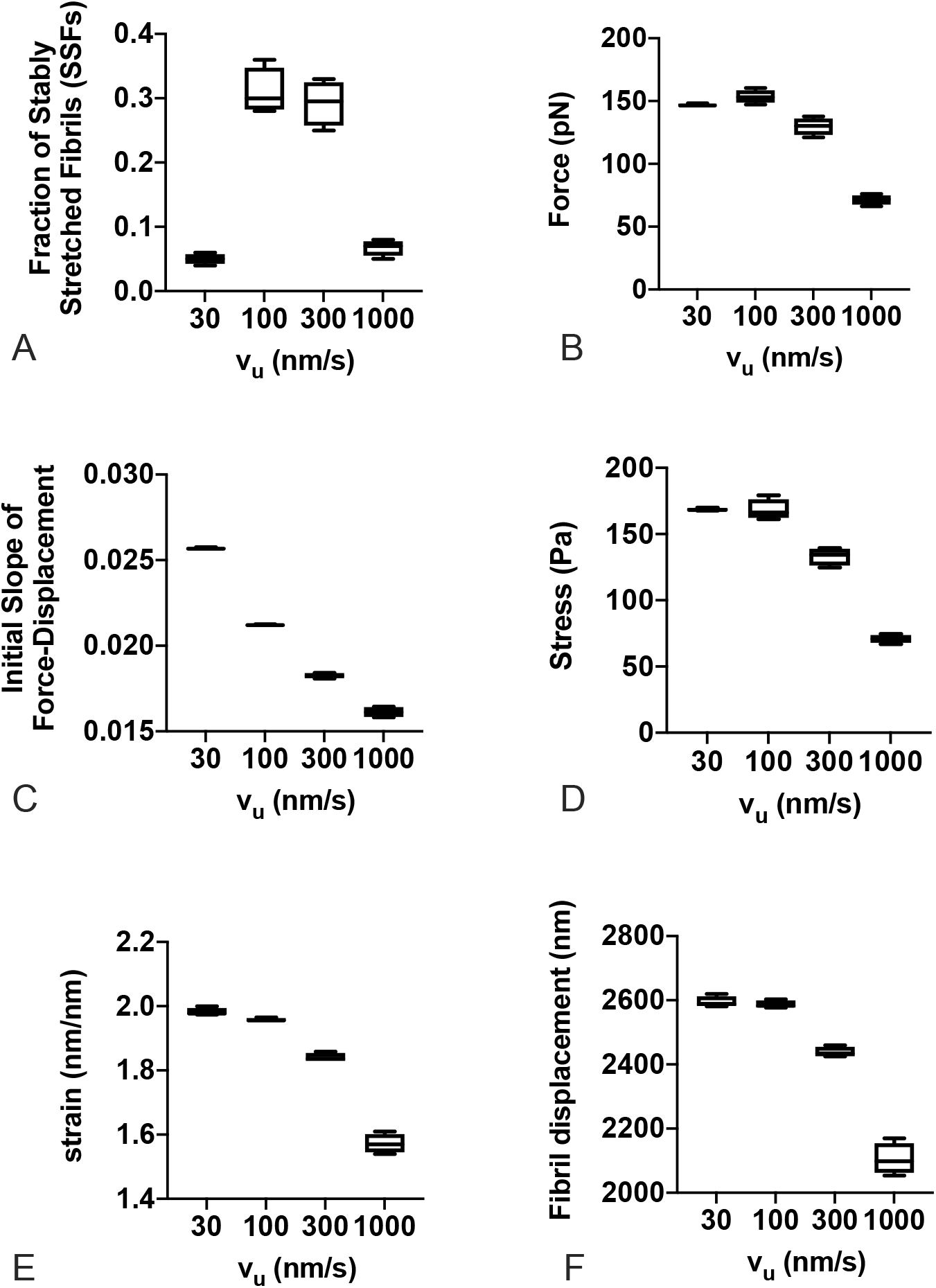
FN fibril morphometry and mechanics as a function of actin velocity. Actin velocity has a pronounced effect on (A) the fraction of SSFs; (B) force transmitted via fibril to the substrate; (C) the initial slope of the force-displacement curve; (D) fibril stress; (E) fibril strain; and (F) fibril displacement. Each condition is the mean of 100 simulations.

## Discussion

In the current work, we have used a previously-developed computational model of FN fibril assembly to investigate the effects of both tissue stiffness and cellular contractile phenotype on fibril growth and mechanics. Our results have found that our model, based on first principles of force-driven FN stretching and assembly, predicts that the morphology and mechanics of FN fibrils depend strongly on the contractile phenotype and only weakly on tissue stiffness. *In silico* experiments demonstrate that as contractility increases, assembled fibrils are smaller, with a shorter length, fewer FN molecules, and a lower fraction of attached integrin clutches. Our results also demonstrate that increasing contractility decreases the stiffness of the fibril. Both stress and strain within the fibril exhibit a maximal value at intermediate contractility, with both low and high contractile states resulting in lower stresses and strains.

We have also examined how tissue stiffness and contractility alter the proportion of two previously identified FN fibril phenotypes: SSFs and FSFs. Our data demonstrates that the proportion of fibrils in each phenotype depends on both tissue stiffness and contractility. At high or low contractility, FSFs are the predominant phenotype. At intermediate contractility, the percentage of SSFs increases with increasing tissue stiffness.

Finally, we have examined the effects of cellular contractile phenotype on the re-stretch of previously assembled FN fibrils. Results indicate that the unloaded actin velocity of the re-stretching cell affects the fibrils: a larger contractile phenotype results in fibrils that are less stretched and have a lower steady-state force magnitude.

One significant impact of the current work is that it provides a mechanistic explanation of two counter-intuitive experimental observations. First, previous studies have shown a bi-phasic response of FN fibril assembly to contractile force magnitude: knocking out traction forces with either myosin inhibitors or actin disrupters results in no FN fibril assembly [8]; but, increasing contractile force also leads to a reduction in FN fibril assembly [23]. Similarly, both cells that generate weak contractile forces, such as epithelial cells, and cells that generate large contractile forces, such as smooth muscle cells, exhibit minimal assembly of FN fibrils [9, 24]. Consistent with this, our model shows no FN fibril assembly at a non-contractile state, that is, v_*u*_ = 0 (data not shown), and shows that as cell contractility increases in the model, cells assemble fibrils that are smaller and shorter.

Given that cellular traction forces increase with substrate stiffness [25, 26], and cellular traction forces drive FN fibril assembly [9], another experimental finding that is perhaps counter-intuitive is that FN fibrillogenesis does not depend strongly on substrate stiffness [27]. Our data again support this experimental finding, demonstrating that substrate stiffness only weakly affects FN morphology and mechanics. Our model suggests that the viscoelasticity of the fibril acts to “buffer” the tissue stiffness; as we have previously described [18], the FN assembly model results in an intermediate mechanics regime in which the presence of the fibril makes a soft surface “appear” stiffer, and a stiff surface “appear” softer. As such, assembly of FN fibrils may serve as a mechanism for mesenchymal cells to adjust and tailor their response to the surrounding tissue mechanics.

While the current work recapitulates several key experimental findings, it fails to accurately predict certain aspects of the FN/force/stiffness relationship. For example, while we have previously demonstrated that FN assembly is not dramatically affected by tissue stiffness, we and others have shown that traction forces still increase in response to increasing tissue stiffness when FN fibril assembly occurs [25, 27]. However, our model predicts that force transmitted via the fibril does not depend on tissue stiffness at all. One possible explanation is that we are predicting the mechanics and morphology of a single FN fibril. A cell attached to a surface will have tens to hundreds of attachments, and thus tens to hundreds of FN fibrils. Our model doesn’t account for or predict the effects of multiple fibrils; thus, it is possible that in the described experimental work, each FN fibril transmits a similar magnitude of traction force, but that increased tissue stiffness results in more fibrils, and thus a larger total force applied to the substrate.

Of particular interest is the role that the two identified FN fibril phenotypes play in mechanotransduction, and how these phenotypes are affected by contractility and tissue stiffness. One could imagine that the cellular response to an FSF, in which the strain and force transmitted to the attached cell are fluctuating randomly, and an SSF, in which the fibril is tonically stretched and a constant force is transmitted, could be dramatically different. Our findings that: SSFs are only assembled in substantial number at an intermediate contractility; re-stretched fibrils exhibit SSF behavior at an intermediate contractile phenotype; and the fraction of SSFs increases with increasing stiffness, suggest that FN fibrils may be optimally tuned to provide mechanical cues at a given contractile state. Cells that can’t maintain this state, either by generating less force or by generating more force, don’t assemble SSFs, and thus may lose a key mechanism to sense tissue stiffness. Investigation of FN fibril mechanics and phenotype, both experimentally and computationally, represents an active area of future work for our group.

## Conclusion

Using a computational model of fibronectin fibril formation, we demonstrate that actin velocity, and in turn contractile phenotype, have a much more profound effect on fibril formation than tissue stiffness. Our work also demonstrates that both tissue stiffness and actin velocity/contractility affect the ratio of two FN fibril phenotypes. At both low and high contractility, cells predominantly assemble fluctuating stretched fibrils (FSFs), while at intermediate contractility, cells assemble more stably stretched fibrils (SSFs), with the percentage of SSFs increasing with tissue stiffness. Together, these results give new insights into how tissue stiffness and cell contractility govern the assembly of FN fibrils.

Understanding the complex relationship between substrate stiffness, contractility, and FN assembly is key to understanding states where FN assembly is desirable, such as wound healing and tissue engineering applications, as well as understanding conditions where FN assembly is not desirable, such as the desmoplastic extracellular matrix of a growing tumor, the fibrotic response to implanted material within the body, and scar tissue in fibrotic disease.

## Acknowledgments

This work was supported by the National Institutes of Health via grant award R01GM115678 (SHW, CAL) and R01GM122855 (SHW, CAL).

